# A replicable and modular benchmark for long-read transcript quantification methods

**DOI:** 10.1101/2024.07.30.605821

**Authors:** Zahra Zare Jousheghani, Noor Pratap Singh, Rob Patro

## Abstract

We provide a replicable benchmark for long-read transcript quantification, and evaluate the performance of some recently-introduced tools on several synthetic long-read RNA-seq datasets. This benchmark is designed to allow the results to be easily replicated by other researchers, and the structure of the underlying `Snakemake` workflow is modular to make the addition of new tools or new data sets relatively easy. In analyzing previously assessed simulations, we find discrepancies with recently-published results. We also demonstrate that the robustness of certain approaches hinge critically on the quality and “cleanness” of the simulated data.

**Availability:** The Snakemake scripts for the benchmark are available at https://github.com/COMBINE-lab/lr_quant_benchmarks, the data used as input for the benchmarks (reference sequences, annotations, and simulated reads) are available at https://doi.org/10.5281/zenodo.13130623.

## 1. Introduction

The sequencing of RNA with long-read sequencing technologies, specifically those from Oxford Nanopore (ONT) and Pacific Biosciences (PacBio), is becoming increasingly common. In recent years, both the throughput (i.e. number of sequenced reads) and the quality (i.e. base-level accuracy) of these technologies has increased substantially (1). These data promise a new perspective on RNA biology, and are proving complementary to existing short-read technologies for applications like transcript discovery and quantification.

However, differences in the process of data generation between long-read and short-read RNA sequencing protocols necessitates the development of new tools that are specifically targeted toward processing long-read data.

In this brief note, we focus specifically on developing a benchmark over a set of simulated data to assess the performance of some recently-developed methods for long read transcript quantification. We aim for our benchmark to be “replicable” in the sense of the “ACM terminology” (2) — that is, it is sufficient for other researchers to repeat the analysis and obtain basically equivalent results. Further, we have implemented the benchmarking setup using `Snakemake` (3), and have designed it to be modular, with the intent that expanding the repertoire of evaluated methods should be simple, and thus researchers can potentially build upon this benchmark to explore the “reproducibility” of the performance of different tools. This desire stems, in part, from the fact that we were not able to replicate the results of a recent benchmark, some of whose data we incorporate herein (4). That result may be repeatable, but it appears neither replicable nor reproducible with just the information currently available.

In evaluating different tools on the benchmarks introduced here, we observe how the performance of different methods can depend, sometimes critically, on the quality and “cleanness” of the underlying simulation, and we also make note of some substantial discrepancies with previous published results.

## 2. Benchmarked tools and data

We evaluate 4 methods across a series of 4 different simulated datasets. Specifically, we evaluate `bambu` (5) (v3.4.1, installed via BioConductor (6) 3.18), `lr-kallisto` (4) (v0.51.0 compiled from source, accompanied by `bustools` (7) v0.43.2 compiled from source), `NanoCount` (8) (v1.1.0 installed from source), and `oarfish` (9) (v0.4.0 and v0.5.0 installed from source). We consider two different configurations of `NanoCount`, since we find that the default filters, while potentially useful when working with experimental data, tend to negatively affect the performance of the method in assessment on simulated data. Likewise, we consider both the previously–released version of `oarfish` (v0.4.0), and v0.5.0 (released concurrently with this manuscript).

### Tools evaluated

While there are many different tools for long-read quantification, and new ones are emerging constantly, we have selected these tools as the initial ones to demonstrate in the replicable benchmark because (a) they all place a substantial focus on *transcript quantification* and not just *transcript discovery* — in fact, `bambu` is the only of these tools that incorporates its own method for transcript discovery (which we disabled for our comparison) and (b) these methods are relatively recent and represent approaches that attempt to resolve multimapping-reads, which persist even in long-read RNA-seq data.

Despite the similarities, there are also substantial differences in the way these methods work. For example, `bambu` operates on spliced-alignments made against the genome and uses an approach that classifies alignments into different categories prior to quantification. Both `NanoCount` and `oarfish` evaluate unspliced alignments made against the transcriptome, and `NanoCount` develops a set of several filters that can be used to remove low-quality alignments from consideration prior to quantification. We note that `NanoCount` was originally developed for processing ONT direct-RNA data, and while recent version provide experimental support for other protocols (mostly by disabling filtering of alignments by orientation), such results should be considered in the context of the experimental support for these other protocols. `Oarfish` implements the filtering capabilities developed in `NanoCount`, and also makes several modifications to the probabilistic model underlying quantification. Finally, `lr-kallisto` modifies pseudoalignment for application to long-read technologies, and performs pseudoalignment of the reads against a target transcriptome; it also modifies the notion of transcript effective length for use with long-read quantification.

### Datasets included

We evaluate the above methods on 4 datasets. Two of these datasets are generated according to the methodology proposed by Ji and Pertea (10). Specifically, these datasets are generated with the `NanoSim` (11) simulator by learning the model parameters from experimental ONT sequencing datasets (both direct-RNA and 1D-cDNA) from Workman et al. (12). We evaluate one simulation of a direct RNA (dRNA) protocol dataset and a second simulation of a cDNA protocol dataset. Further, we follow the methodology of Ji and Pertea where abundances are seeded using quantification estimates from an experimental sample, and where simulated reads are drawn only from protein-coding and long noncoding RNA (but where quantification is performed against the entire reference transcript catalog). Following the protocol of Ji and Pertea, these experiments are quantified against a RefSeq (13) annotation of the human reference genome and transcriptome (GRCh38.p14). These simulations pose a more challenging quantification task than the other two datasets described next. This is driven by several factors, including the (empirically derived) inclusion of “background” reads in the simulation, and higher error rates (though all model parameters are learned by `NanoSim` from the experimental data). We refer to these datasets as `ts-dRNA` and `ts-cDNA`, respectively, since the simulation follows the protocol that was originally introduced in the `TranSigner` paper (10).

The next two datasets we consider were originally introduced in the `IsoQuant` paper (14) and recently used in the `lr-kallisto` paper (4). These simulated data consist of one ONT and one PacBio dataset, both of which are quantified against the LRGASP (15) mouse annotation using the `GRCm39-based mouse`^1^ genome and `GENCODE VM27-based mouse annotation`^2^. The PacBio dataset was simulated with `PBSim` (16) with an error rate of 1.4% (further details in (14)) and the (cDNA) ONT dataset was simulated with `NanoSim` (11) with an error rate of 2.8%. Of note, the ONT dataset was simulated by disabling the default `NanoSim` behavior of creating background “unaligned” reads, which results in a “cleaner” simulation then when such reads are included in the simulated data. In the context of short-read RNA-seq simulation, for example, it has previously been demonstrated (17) that the inclusion of such “background” reads adds an important aspect of realism to the simulation which can affect the overall and even relative performance of different methods. We refer to these datasets as `iq-PacBio` and `iq-ONT`, respectively, since these simulations were first introduced in the `IsoQuant` (14) paper, and were among the simulations later used in the `lr-kallisto` (4) paper.

## 3. Results

We compute several different metrics to assess the evaluated methods, including the mean absolute relative difference (MARD) between (raw) true and predicted counts, the Pearson correlation of the (log-transformed) predicted and true counts, the Spearman-*ρ* and Kendall-*τ* correlation coefficients between the predicted and true counts, the Concordance correlation coefficient (CCC; computed on logtransformed counts), and the normalized root mean squared error (NRMSE; over the raw counts and using the variant where the RMSE is normalized by the mean of the observed values). Finally, to comport with the manner in which quantification tools are most-often used, and to ensure fairness and avoid unnecessary bias in the evaluation, we calculate and report transcriptome-wide metrics for all methods.

### Evaluation on TranSigner-protocol simulated data

We observe that, in general, all methods, save `NanoCount`, perform slightly better on the `ts-dRNA` dataset (Table 1) than the `ts-cDNA` dataset (Table 2). Nonetheless, despite a modest shift across computed metrics, the relative performance of the methods is quite consistent between these datasets. Specifically, `oarfish` 0.5 with the coverage model enabled offers the best performance under most metrics, followed by `oarfish` 0.5 and 0.4 without the coverage model. After these approaches, `bambu` demonstrates the next best performance across a variety of metrics, followed by `oarfish` 0.4 with the coverage model enabled. We also observed that across the different metrics, for both the `ts-cDNA` and `ts-dRNA` simulations, `lr-kallisto` did not perform as well as these other methods, while it exhibited superior performance compared to the evaluated variations of `NanoCount`.

**Table 1.**
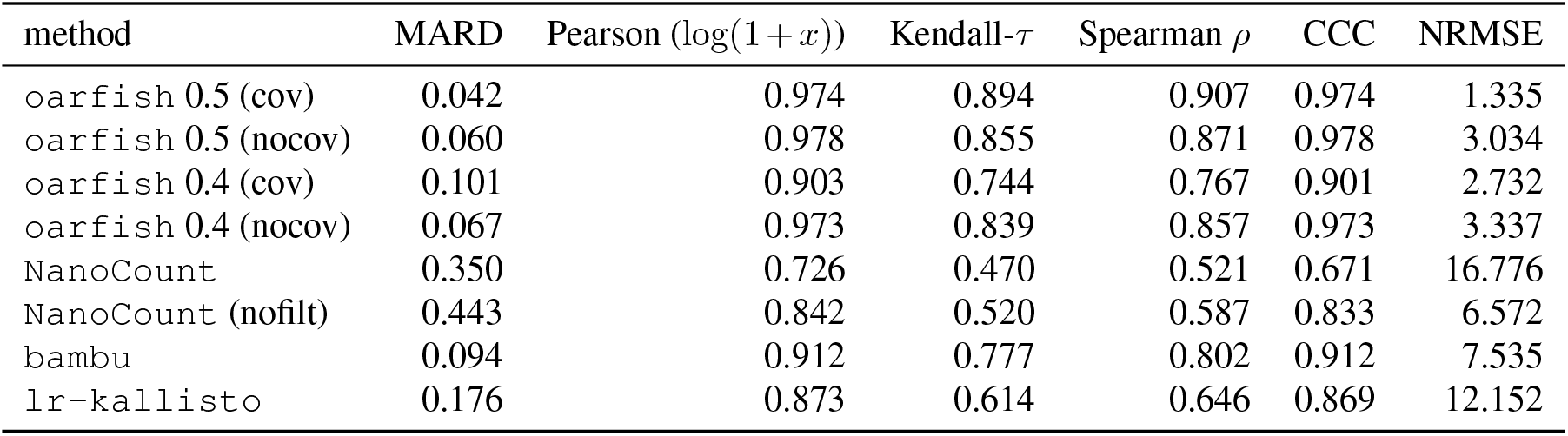
Evaluation metrics on the ts-dRNA simulation for the evaluated methods.

| method | MARD | Pearson ($\log(1+x)$) | Kendall-τ | Spearman $\rho$ | CCC | NRMSE |
| --- | --- | --- | --- | --- | --- | --- |
| oarfish 0.5 (cov) | 0.042 | 0.974 | 0.894 | 0.907 | 0.974 | 1.335 |
| oarfish 0.5 (nocov) | 0.060 | 0.978 | 0.855 | 0.871 | 0.978 | 3.034 |
| oarfish 0.4 (cov) | 0.101 | 0.903 | 0.744 | 0.767 | 0.901 | 2.732 |
| oarfish 0.4 (nocov) | 0.067 | 0.973 | 0.839 | 0.857 | 0.973 | 3.337 |
| NanoCount | 0.350 | 0.726 | 0.470 | 0.521 | 0.671 | 16.776 |
| NanoCount (nofilt) | 0.443 | 0.842 | 0.520 | 0.587 | 0.833 | 6.572 |
| bambu | 0.094 | 0.912 | 0.777 | 0.802 | 0.912 | 7.535 |
| lr-kallisto | 0.176 | 0.873 | 0.614 | 0.646 | 0.869 | 12.152 |

**Table 2.** Evaluation metrics on the ts-cDNA simulation for the evaluated methods.

| method | MARD | Pearson ($\log(1+x)$) | Kendall-τ | Spearman $\rho$ | CCC | NRMSE |
| --- | --- | --- | --- | --- | --- | --- |
| oarfish 0.5 (cov) | 0.065 | 0.951 | 0.850 | 0.869 | 0.950 | 1.865 |
| oarfish 0.5 (nocov) | 0.087 | 0.962 | 0.816 | 0.839 | 0.961 | 4.178 |
| oarfish 0.4 (cov) | 0.136 | 0.867 | 0.707 | 0.739 | 0.865 | 3.204 |
| oarfish 0.4 (nocov) | 0.092 | 0.954 | 0.805 | 0.829 | 0.953 | 4.703 |
| NanoCount | 0.327 | 0.704 | 0.479 | 0.529 | 0.591 | 21.549 |
| NanoCount (nofilt) | 0.435 | 0.889 | 0.561 | 0.632 | 0.885 | 6.731 |
| bambu | 0.118 | 0.903 | 0.755 | 0.786 | 0.903 | 7.867 |
| lr-kallisto | 0.197 | 0.871 | 0.612 | 0.649 | 0.864 | 12.246 |

In addition to the numerical results provided in Tables 1 and 2, Figures 1 and 2 provide density plots of the logtransformed quantification estimates from different tools compared to the ground truth counts. The density plots for the `ts-dRNA` and `ts-cDNA` datasets, depicted in Figures 1 and 2, respectively, illustrate a strong correlation for most methods, and notably less variation around the *y* = *x* diagonal for most `oarfish` variants.

**Fig. 1.**
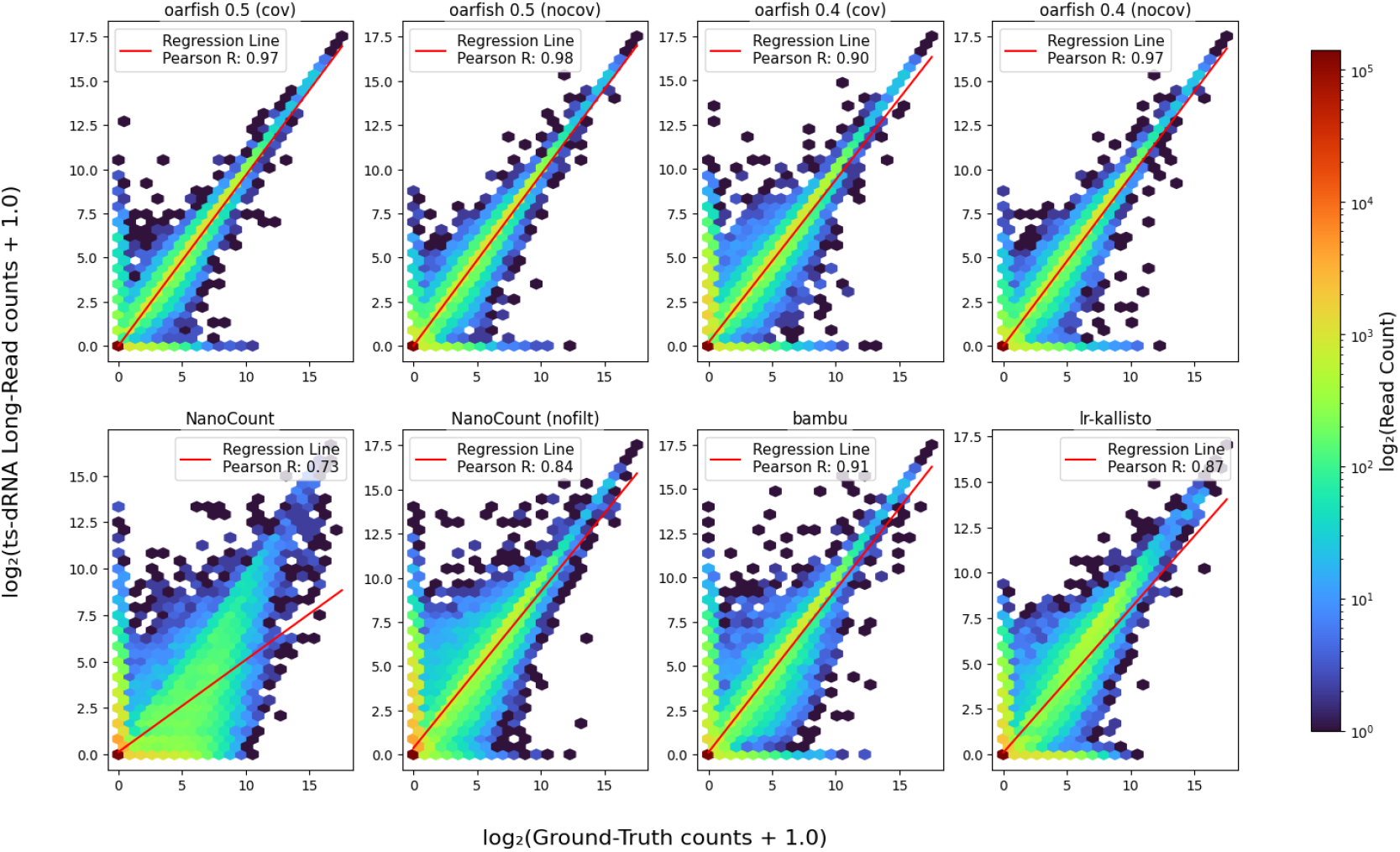
Density plot of log values of different quantification tools and the log value of the ground truth for `ts-dRNA` dataset.

**Fig. 2.**
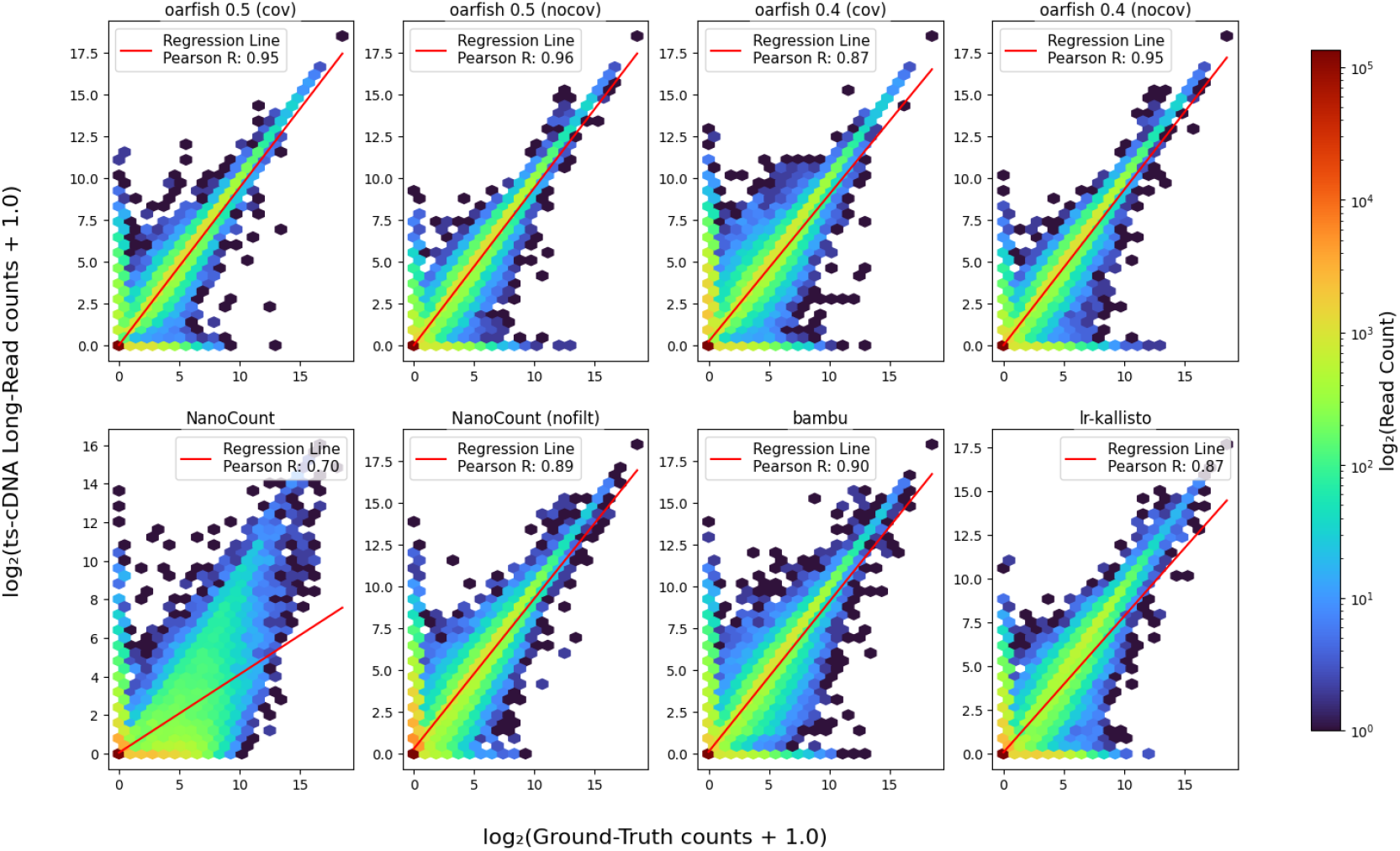
Density plot of log values of different quantification tools and the log value of the ground truth for `ts-cDNA` dataset.

Likewise, Fig. S1 presents plots comparing each of the Pearson correlation, CCC, Kendall-*τ*, NRMSE, and MARD metrics against the Spearman-*ρ* correlation coefficient. To simplify the presentation, we use the inverse values of NRMSE and MARD (i.e., NRMSE^*−*1^ and MARD^*−*1^) on the plots. In these plots, better performance on each pairwise combination of metrics is indicated by the corresponding marks being closer to the top-right corner.

We observe a notable improvement in the performance of `NanoCount`, particularly on the direct-RNA dataset, when its filters are turned off. However, its performance across the different metrics is still relatively lower compared to the other methods. This is likely due to, in part, to its much less stringent termination condition for the EM procedure than all of the other methods — by default it chooses a much larger “early stopping” threshold for the allowable change in transcript abundances between subsequent rounds of the EM algorithm than the other tools, and it runs for fewer than 20 iterations on these samples while most of the other methods run for many dozens or hundreds of iterations. The termination criterion for `NanoCount` can be adjusted from the command line, but we did not attempt to optimize it further in these benchmarks.

### Evaluation on IsoQuant-protocol simulated data

For the `IsoQuant`-protocol simulations — on both the ONT (`iq-ONT`) and PacBio (`iq-PacBio`) data — we observe that all methods exhibit a good performance across most of the computed metrics. This could be due to the substantially “cleaner” data, which leads to all methods allocating a very high fraction of the simulated reads to the reference transcriptome. The performance metrics we record are reported in Table 3 for the `iq-ONT` simulation and in Table 4 for the `iq-PacBio` data. As depicted in Fig. S2, the relative performance of the different methods is similar to that observed on the `TranSigner`-protocol simulations. Still, the differences between methods are vastly reduced, and most tested methods perform similarly on these data (though `NanoCount` still trails behind the other methods somewhat). These quantitative results are also observed in the density plots for the `iq-PacBio` and `iq-ONT` datasets (Figures 3 and 4), which indicate a similar Pearson correlation of log-transformed counts between the quantification results from most different tools and the ground truth. For the `iq-PacBio` dataset, both configurations of `NanoCount` exhibit a lower correlation and greater variation around the *y* = *x* diagonal. Yet, in absolute terms, all methods tended to perform better in the `iq-PacBio` simulation than the `iq-ONT` simulation. For example, the `iq-PacBio` simulation shows very high performance across computed metrics for most methods, owing, presumably, to the combination of the low error rate (1.2%), the lack of any simulated “noise”, and potentially less multimapping of the simulated reads. In this simulation, `oarfish` and `NanoCount` quantifications allocate more than 99.5% of the simulated reads, `bambu` allocates 96.3%, and the more-stringent pseudoalignment approach of `lr-kallisto` allocates 93.4% of the simulated reads.

**Table 3.** Evaluation metrics on the iq-ONT simulation for the evaluated methods.

| method | MARD | Pearson ($\log(1+x)$) | Kendall-τ | Spearman $\rho$ | CCC | NRMSE |
| --- | --- | --- | --- | --- | --- | --- |
| oarfish 0.5 (cov) | 0.062 | 0.960 | 0.926 | 0.953 | 0.960 | 2.146 |
| oarfish 0.5 (nocov) | 0.061 | 0.962 | 0.932 | 0.959 | 0.961 | 2.109 |
| oarfish 0.4 (cov) | 0.086 | 0.960 | 0.884 | 0.912 | 0.960 | 2.260 |
| oarfish 0.4 (nocov) | 0.061 | 0.959 | 0.928 | 0.955 | 0.958 | 2.120 |
| NanoCount | 0.270 | 0.902 | 0.731 | 0.789 | 0.897 | 4.798 |
| NanoCount (nofilt) | 0.298 | 0.943 | 0.727 | 0.798 | 0.943 | 3.306 |
| bambu | 0.090 | 0.936 | 0.883 | 0.919 | 0.936 | 3.971 |
| lr-kallisto | 0.094 | 0.953 | 0.885 | 0.920 | 0.953 | 2.090 |

**Table 4.** Evaluation metrics on the iq-PacBio simulation for the evaluated methods.

| method | MARD | Pearson ($\log(1+x)$) | Kendall-τ | Spearman $\rho$ | CCC | NRMSE |
| --- | --- | --- | --- | --- | --- | --- |
| oarfish 0.5 (cov) | 0.013 | 0.986 | 0.975 | 0.980 | 0.986 | 0.984 |
| oarfish 0.5 (nocov) | 0.015 | 0.984 | 0.971 | 0.977 | 0.984 | 1.130 |
| oarfish 0.4 (cov) | 0.015 | 0.986 | 0.973 | 0.979 | 0.985 | 0.978 |
| oarfish 0.4 (nocov) | 0.018 | 0.980 | 0.966 | 0.973 | 0.980 | 1.314 |
| NanoCount | 0.054 | 0.957 | 0.936 | 0.963 | 0.957 | 3.307 |
| NanoCount (nofilt) | 0.075 | 0.934 | 0.921 | 0.957 | 0.934 | 3.507 |
| bambu | 0.019 | 0.983 | 0.969 | 0.977 | 0.983 | 3.262 |
| lr-kallisto | 0.031 | 0.990 | 0.965 | 0.977 | 0.989 | 3.169 |

**Fig. 3.**
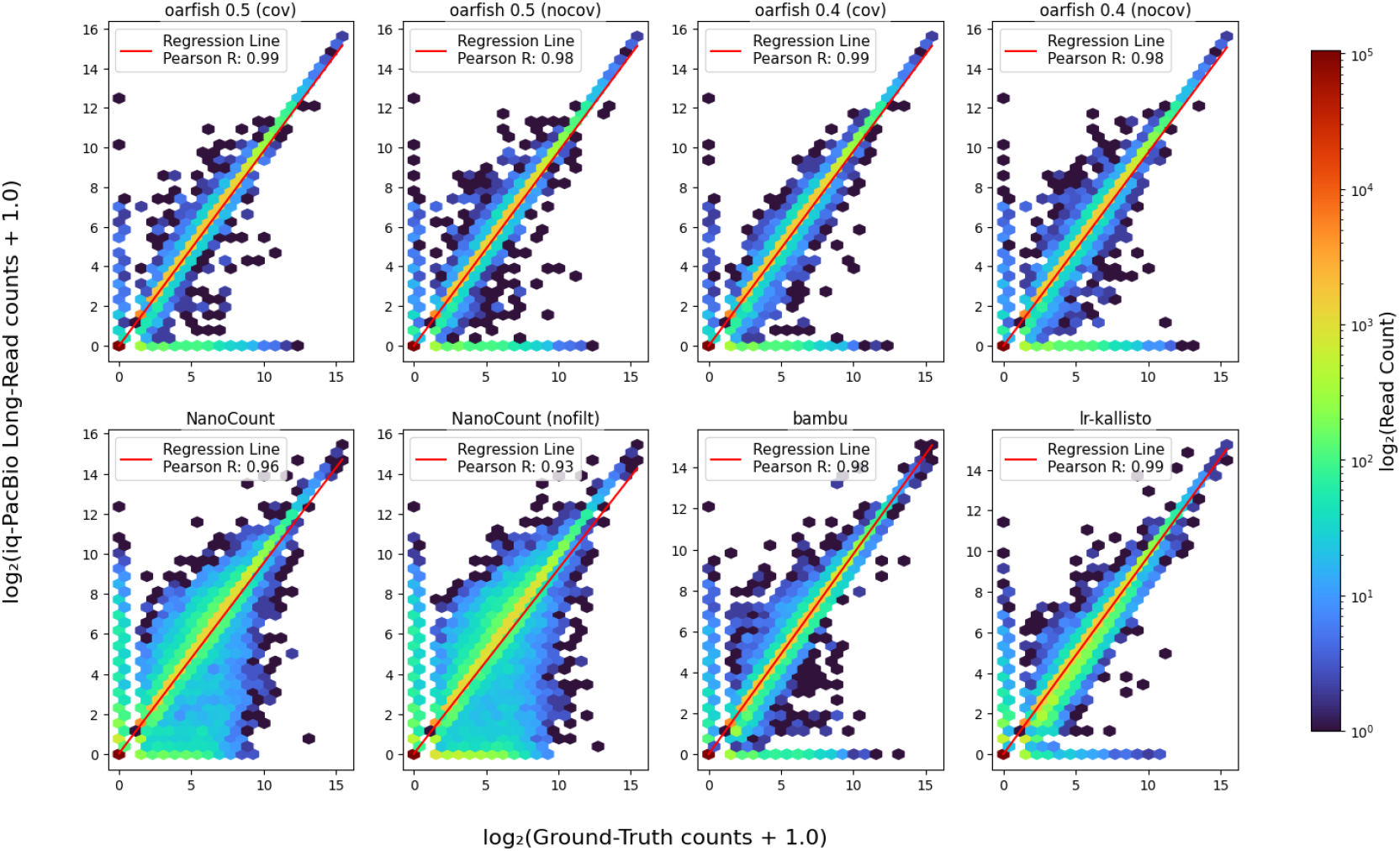
Density plot of log values of different quantification tools and the log value of the ground truth for `iq-PacBio` dataset.

**Fig. 4.**
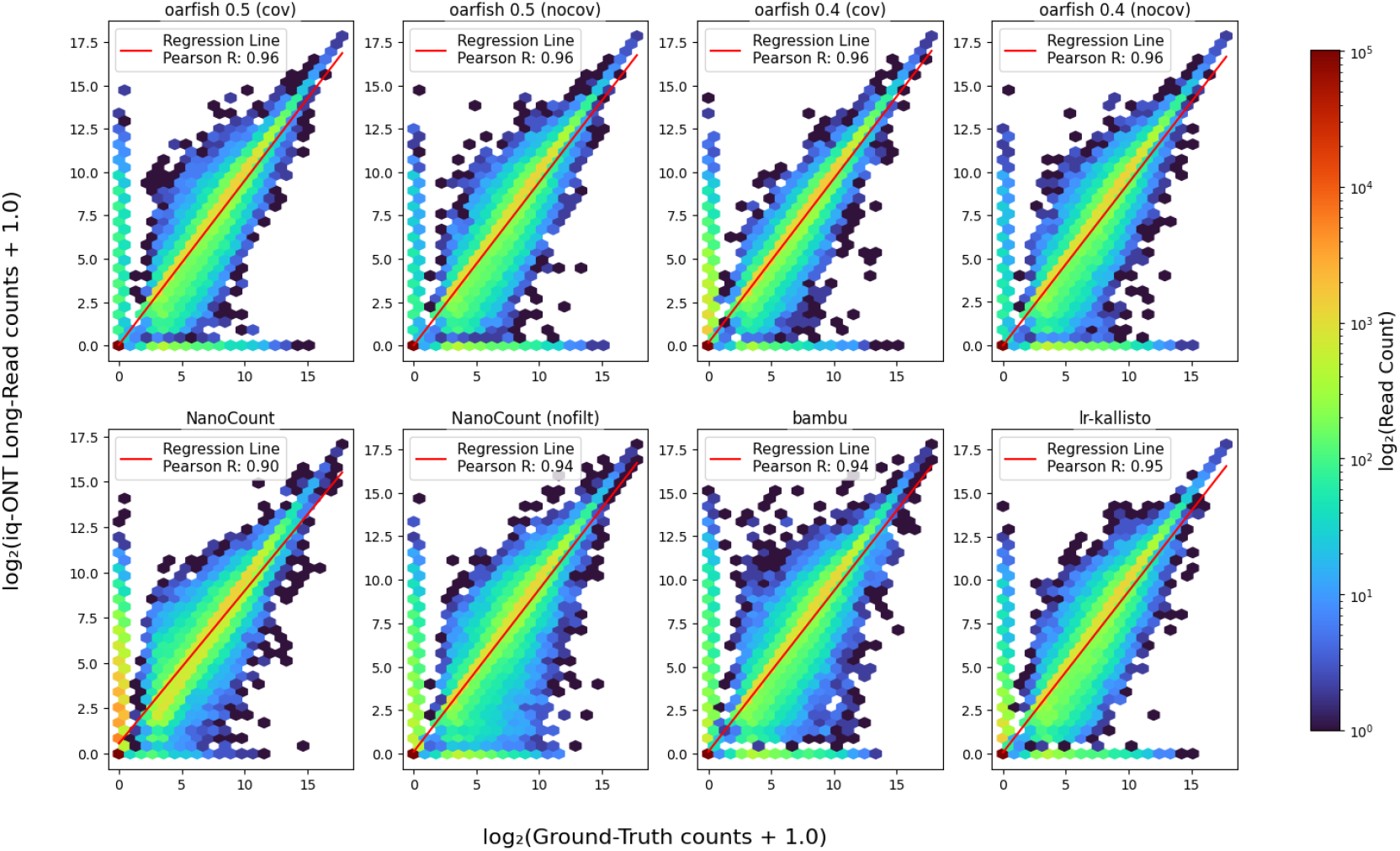
Density plot of log values of different quantification tools and the log value of the ground truth for `iq-ONT` dataset.

However, we note a substantial discrepancy between the performance reported for `oarfish` by Loving et al. (4) and what is observed here. Specifically, while the newer version of `oarfish` (0.5) generally performs best overall, especially when the coverage model is enabled, the previous version of `oarfish` (0.4) — presumably the one benchmarked by Loving et al. (4) (though no explicit version was mentioned in that manuscript) — exhibits a performance that is similar to the other methods. It is, therefore, unclear, how the results arrived at by Loving et al. (4) on this data, which suggest the substantial under-performance of `oarfish` compared to `lr-kallisto` and `bambu`, were obtained.

Specifically, Loving et al. report that `lr-kallisto` outperforms other methods on the `iq-ONT` and `iq-PacBio` datasets based on several metrics — including Spearman-*ρ* correlation, Pearson correlation, CCC, and NRMSE — which does not concord with the results we observe here on these same datasets. While the specific values of certain metrics can vary based on decisions such as normalization (e.g. how the RMSE is normalized in the NRMSE metric) or count transformation (e.g. is the metric computed on raw or logtransformed counts), others, such as the Spearman-*ρ* Depend on the relative ranking of the values and are therefore invariant to monotonic transformations and hence comparable.

On the `iq-ONT` simulation, Loving et al. note a Spearman correlation for `lr-kallisto` of 0.92 and report a corresponding value of 0.7 for `oarfish`. Yet, the results here, on the same simulated dataset (shown in Table 3), indicate that `oarfish` (even version 0.4) performs very similarly to `lr-kallisto` under these metrics (Spearman-*ρ* = 0.91).

Likewise, we also observe that `bambu` obtains a Spearman-*ρ* of 0.92 with the true counts on this data, and thus performs more closely to `lr-kallisto` than is reported by Loving et al. The same trend is observed for the `iq-PacBio` dataset, where Loving et al. report `lr-kallisto` outperforming other tools with a Spearman-*ρ* of 0.97, and specifically outperforming `oarfish` for which they report a Spearman-*ρ* of 0.89. However, the results in Table 4 show that `oarfish` (v0.4) performs very similarly to `lr-kallisto` in this simulation (Spearman-*ρ* = 0.98), and that, again, the performance between `lr-kallisto` and `bambu` appears closer than reported in that manuscript (wherein `bambu` is reported to have a Spearman-*ρ* correlation of 0.96 whereas here we observe 0.98). Similar differences can also be observed with respect to other computed metrics.

## 4. Conclusions

We have introduced a replicable benchmark for long-read transcript quantification methods. The goal of this benchmark is to allow users to easily replicate the results we observe here, as well as to add new tools and potentially new data to the benchmark suite to compare their performance against the core set of tools that we have currently included.

Additionally, this work highlights the difficulty in replicating the results claimed by Loving et al., and the substantially different performance observed for specific tools, even when evaluating the same simulated data.

The results here also demonstrate that the tested methods display a wide spectrum of robustness to data quality and cleanness. The benchmark presented in this work, using the simulation protocol from Ji and Pertea (10), suggests that as the simulation setup changes, the absolute performance of different quantification tools changes considerably (e.g. `lr-kallisto` goes from performing similarly to other methods to considerably under-performing them). Though these specific higher error-rate simulations are different than those described in Loving et al. (4) — wherein it is claimed that `lr-kallisto` performs similar to and often better than, several other approaches, even as the error rate increases — the results are qualitatively at odds with the description in that work. This is true with respect not only to the performance of `oarfish`, but also that of `bambu`, which performs better in this benchmark than in that of Loving et al. Nonetheless, these observations indicate that further investigation may be warranted as to whether or not the currently-proposed modifications of pseudoalignment are adequate to robustly quantify long-read sequencing data under more realistic scenarios, as opposed to methods operating on the output of tools such as `minimap2` (18), that have already proven to be well-tested and robust when aligning long-read RNA-seq data.

Of course, an important caveat to issue for all assessments on simulated data is that these data rely on simplified and idealized data generation procedures, and tend to lack certain, sometimes important, characteristics of experimental data. As a result, they may paint an overly-optimistic picture of the potential performance of different tools. Therefore, in the broader context of tool and method assessment, it is important to complement analyses on simulated data with analyses on experimental data. Yet, such assessments often have their own caveats related to the existence of a known ground truth or to the limited complexity of the data subset on which known ground truth results can be easily obtained (e.g. spikein and sequin (19) experiments). Nonetheless, evaluations on simulated data, as carried out here, are an important aspect of method assessment, and we hope that the community finds the benchmarks presented here useful and builds upon them.

## S1. Software versions for alignment and quantification tools

- `bambu`: `v3.4.1` (in `R` v4.3.3)
- `bustools`: `v0.43.2`
- `kallisto`: `v0.51.0`^3^.
- `minimap2`: `2.28-r1209`
- `NanoCount`: `v1.1.0`^4^
- `oarfish`: `v0.4.0`^5^; `v0.5.0`^6^.

## S2. Notes with respect to replicating IsoQuant-protocol benchmarks

While we have attempted, where possible, to use the same parameters provided in the scripts of Loving et al. for running the `lr-kallisto` tool, we encountered certain difficulties, and also observed some limitations as well as some inconsistencies in the way certain metrics were computed, which we note below.

- The `--threshold` parameter cannot be set, as the logical condition to check for a valid threshold (i.e. > 0 and < 1) uses a chained comparison, which is not supported in `C++`. Thus, the check always fails, and the threshold is set to the default of 0.8 (with a message to this effect displayed to the console when running the tool). While this will be a problem for changing the `--threshold` parameter until this issue is resolved, it should not affect the results on the `IsoQuant`-protocol simulations, as the default parameter was used there anyway. Because of this limitation, we were also unable to modify this parameter for the `TranSigner`-protocol simulations and have kept it at the default value of 0.8.
- On the first computer on which we attempted to run `lr-kallisto`, we were unable to build the index on either of the reference transcriptomes without encountering a segmentation fault. We were able to overcome this issue by moving the evaluations to a different machine, but were unable to diagnose the root cause of the segmentation fault. Others may therefore encounter this issue until it is resolved upstream.
- The `--error-rate` parameter, present in some of the scripts, but which as far as can be discerned was not used in the evaluation of the simulated data sets considered in this manuscript, is not a valid command line parameter in the software, and the relevant related code has been commented out. Thus, this parameter had to be removed from invocations of `lr-kallisto`.
- In the manuscript and accompanying scripts of (4), it was not clear which versions of different tools were used. We have therefore used the previously-released version of `oarfish` (v0.4.0; released on March 29, 2024), and `bambu` v3.4.1 (which is part of the BioConductor 3.18 release from October 25, 2023).
- We had initially considered attempting to replicate some of the higher error rate simulations mentioned in the supplementary material of (4). However, the relevant parameters and invocations necessary to simulate PacBio data, while sweeping across various error rates, were not available in the repository^7^. Additionally, the `TranSigner`-protocol simulations that are included in the manuscript already serve to evaluate the tools under higher error rate ONT data.
- In the scripts accompanying (4), reported performance metrics are not always computed in the same manner. For example, the computation of CCC is sometimes carried out on log-transformed counts^8^ and is sometimes carried out on raw counts^9^, but still with an added pseudocount of 1. While there may be motivations for different choices in certain scenarios, we have elected to compute the metrics in a consistent manner throughout, and note, when we introduce each metric, whether it is evaluated on log-transformed or raw counts.

## S3. Notes with respect to evaluating bambu in the TranSigner-protocol simulations

- In the RefSeq reference, we note some small discrepancies in the transcript set present in the available RNA fasta file^10^ and what is present in the available GTF annotation^11^. Specifically, the latter has some extra transcripts, and additional variants of some transcripts present in the former. Since `bambu` quantifies against the genome and annotation, it therefore presents a slightly different set of transcripts in its output. To address this discrepancy, we took any transcript names that collide when removing the portion after the initial ‘.’ character, and summed their assigned read count. Any transcript models that were present only in the GTF annotation and not in the transcriptome `FASTA` file were removed from the `bambu` quantification results prior to evaluation.

## S4. Additional figures for presented performance metrics

**Fig. S1.**
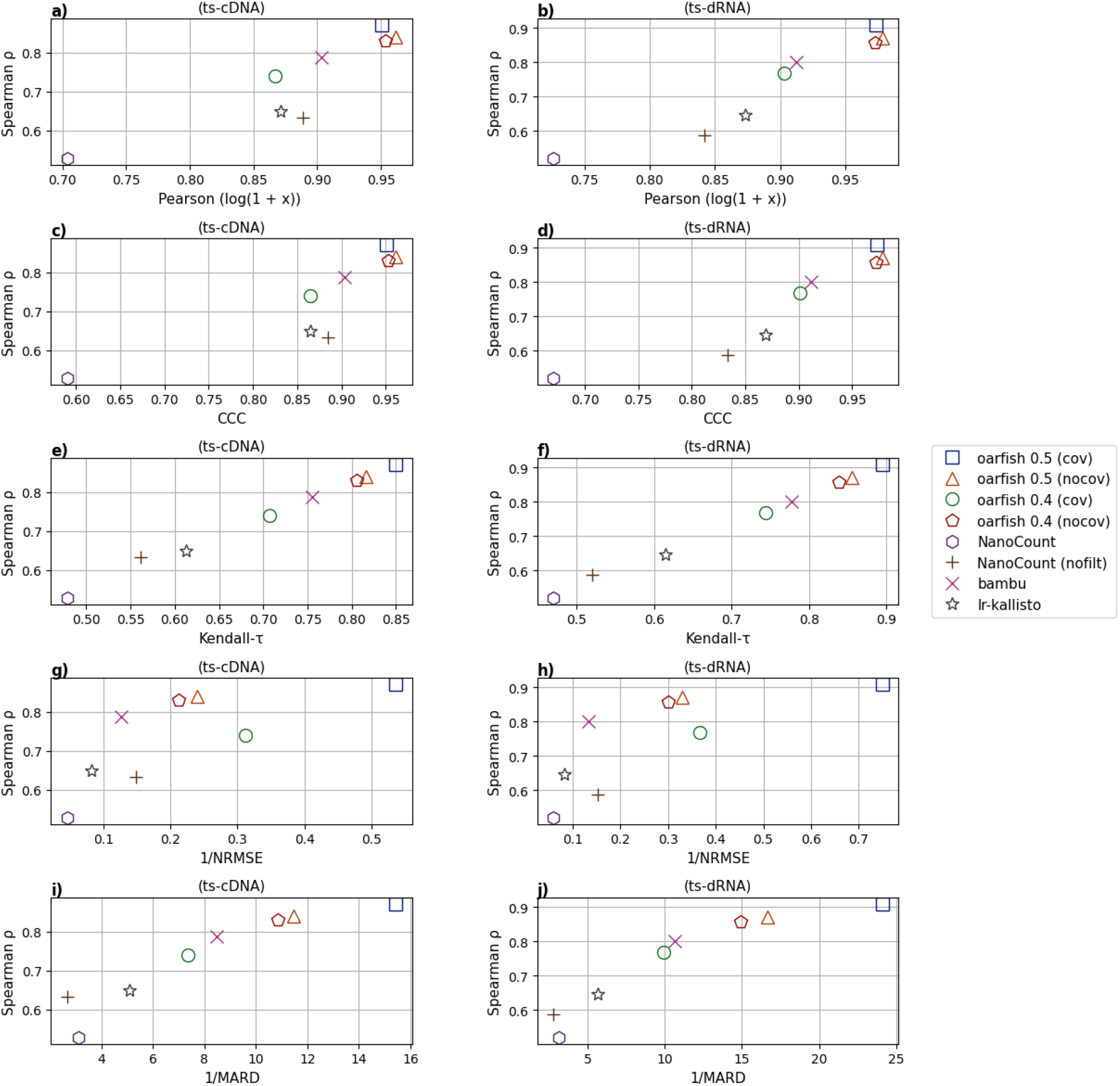
comparison between Spearman-*ρ* correlation and other metrics such as Pearson correlation, CCC, Kendall-*τ*, NRMSE^*−*1^ and MARD^*−*1^, for quantification results obtained from different tools for the `ts-cDNA` and `ts-dRNA` datasets: (a), (b) Spearman correlation compared to Pearson correlation for all tools on `ts-cDNA` and `ts-dRNA` datasets, respectively. (c), (d) Spearman correlation compared to CCC for all tools on `ts-cDNA` and `ts-dRNA` datasets, respectively. (e), (f) Spearman correlation compared to Kendall-*τ* for all tools on `ts-cDNA` and `ts-dRNA` datasets respectively.(g), (h) Spearman correlation compared to NRMSE^*−*1^ for all tools on `ts-cDNA` and `ts-dRNA` datasets, respectively. (i), (j) Spearman correlation compared to MARD^*−*1^ for all tools on `ts-cDNA` and `ts-dRNA` datasets, respectively.

**Fig. S2.**
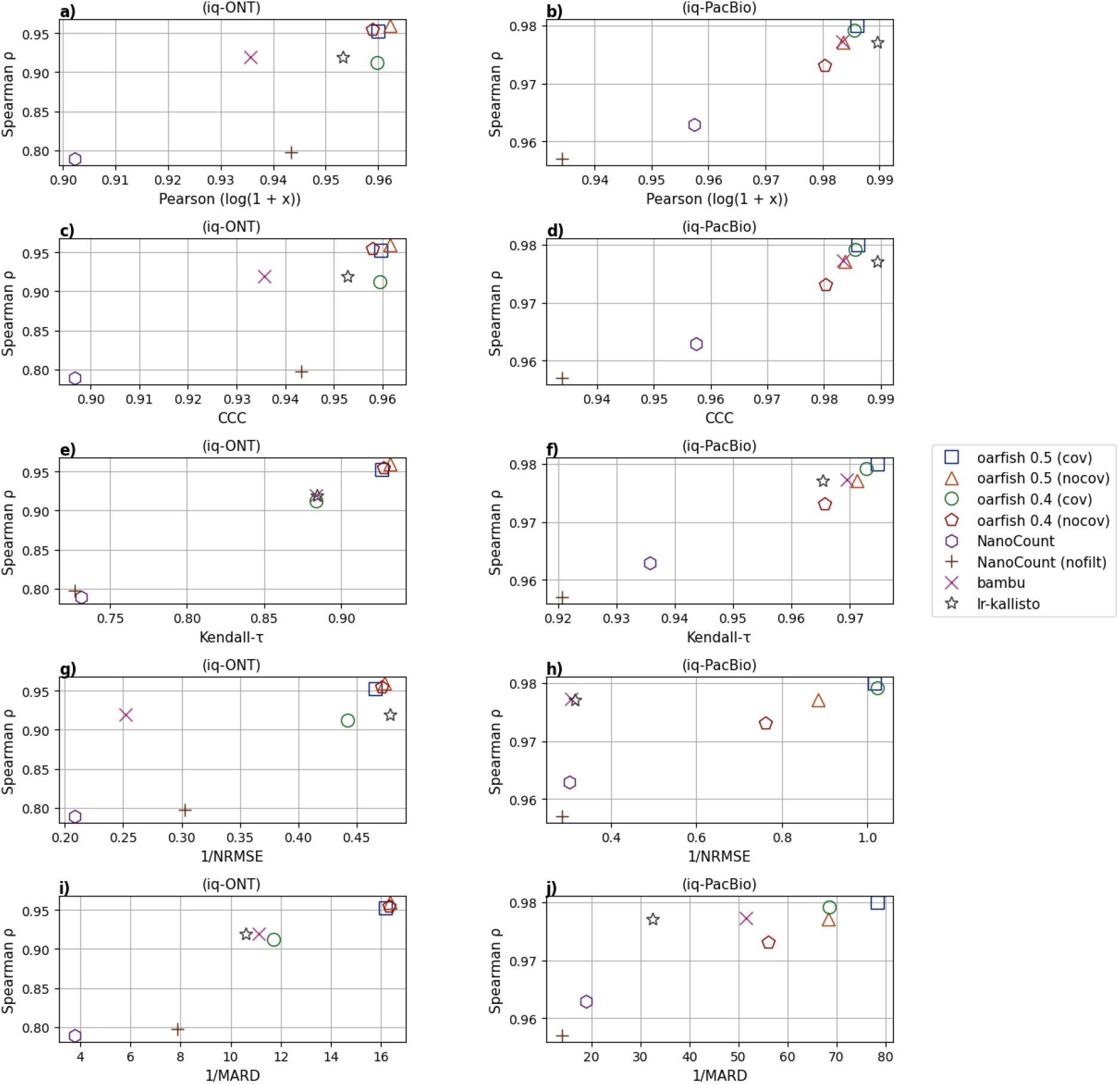
Comparison between Spearman-*ρ* correlation and other metrics such as Pearson correlation, CCC, Kendall-*τ*, NRMSE^*−*1^ and MARD^*−*1^, for quantification results obtained from different tools for the `iq-ONT` and `iq-PacBio` datasets: (a), (b) Spearman correlation compared to Pearson correlation for all tools on `iq-ONT` and `iq-PacBio` datasets, respectively. (c), (d) Spearman correlation compared to CCC for all tools on `iq-ONT` and `iq-PacBio` datasets, respectively. (e), (f) Spearman correlation compared to Kendall-*τ* for all tools on `iq-ONT` and `iq-PacBio` datasets respectively.(g), (h) Spearman correlation compared to NRMSE^*−*1^ for all tools on `iq-ONT` and `iq-PacBio` datasets, respectively. (i), (j) Spearman correlation compared to MARD^*−*1^ for all tools on `iq-ONT` and `iq-PacBio` datasets, respectively.

## Footnotes

1 https://www.synapse.org/#!Synapse:syn25683365

2 https://www.synapse.org/#!Synapse:syn25683629

3 tarball SHA256 sum `efeb0191c1a6a0d6de69111fb66f4bda51ff31fb40c513280f072bd44556f80d`.

4 Version 1.1.0 is not available via pip or bioconda, so it was installed from source using the head of the “master” branch, corresponding to git commit hash `3ae643c6aae87166b56269166c6c8e472b9ec0e7`.

5 https://github.com/COMBINE-lab/oarfish/archive/refs/tags/v0.4.0.tar.gz

6 https://github.com/COMBINE-lab/oarfish/archive/refs/tags/v0.5.0.tar.gz

7 https://github.com/pachterlab/LSRRSRLFKOTWMWMP_2024/blob/main/Figures/SupplementalFigure4/simulate_PB.py

8 https://github.com/pachterlab/LSRRSRLFKOTWMWMP_2024/blob/00399aca1c4e41091867466b0b4924af150a4dd6/Figures/SupplementalFigure1/comparison.py#L118

9 https://github.com/pachterlab/LSRRSRLFKOTWMWMP_2024/blob/00399aca1c4e41091867466b0b4924af150a4dd6/Figures/Figure3/lr-kallisto-Fig3-revised.ipynb

10 https://ftp.ncbi.nlm.nih.gov/genomes/all/annotation_releases/9606/110/GCF_000001405.40_GRCh38.p14/GCF_000001405.40_GRCh38.p14_rna.fna.gz

11 https://ftp.ncbi.nlm.nih.gov/genomes/all/annotation_releases/9606/110/GCF_000001405.40_GRCh38.p14/GCF_000001405.40_GRCh38.p14_genomic.gtf.gz

## Bibliography

1. Francisco J. Pardo-Palacios, Dingjie Wang, Fairlie Reese, Mark Diekhans, Sílvia Carbonell-Sala, Brian Williams, Jane E. Loveland, Maite D. María, Matthew S. Adams, Gabriela Balderrama-Gutierrez, Amit K. Behera, Jose M. Gonzalez Martinez, Toby Hunt, Julien Lagarde, Cindy E. Liang, Haoran Li, Marcus Jerryd Meade, David A. Moraga Amador, Andrey D. Prjibelski, Inanc Birol, Hamed Bostan, Ashley M. Brooks, Muhammed Hasan undefinedelik, Ying Chen, Mei R. M. Du, Colette Felton, Jonathan Göke, Saber Hafezqorani, Ralf Herwig, Hideya Kawaji, Joseph Lee, Jian-Liang Li, Matthias Lienhard, Alla Mikheenko, Dennis Mulligan, Ka Ming Nip, Mihaela Pertea, Matthew E. Ritchie, Andre D. Sim, Alison D. Tang, Yuk Kei Wan, Changqing Wang, Brandon Y. Wong, Chen Yang, If Barnes, Andrew E. Berry, Salvador Capella-Gutierrez, Alyssa Cousineau, Namrita Dhillon, Jose M. Fernandez-Gonzalez, Luis Ferrández-Peral, Natàlia Garcia-Reyero, Stefan Götz, Carles Hernández-Ferrer, Liudmyla Kondratova, Tianyuan Liu, Alessandra Martinez-Martin, Carlos Menor, Jorge Mestre-Tomás, Jonathan M. Mudge, Nedka G. Panayotova, Alejandro Paniagua, Dmitry Repchevsky, Xingjie Ren, Eric Rouchka, Brandon Saint-John, Enrique Sapena, Leon Sheynkman, Melissa Laird Smith, Marie-Marthe Suner, Hazuki Takahashi, Ingrid A. Youngworth, Piero Carninci, Nancy D. Denslow, Roderic Guigó, Margaret E. Hunter, Rene Maehr, Yin Shen, Hagen U. Tilgner, Barbara J. Wold, Christopher Vollmers, Adam Frankish, Kin Fai Au, Gloria M. Sheynkman, Ali Mortazavi, Ana Conesa, and Angela N. Brooks. Systematic assessment of long-read rna-seq methods for transcript identification and quantification. Nature Methods, 21(7):1349–1363, June 2024. ISSN 1548-7105. doi: 10.1038/s41592-024-02298-3.

2. Hans E. Plesser. Reproducibility vs. Replicability: A Brief History of a Confused Terminology. Frontiers in Neuroinformatics, 11, January 2018. ISSN 1662-5196. doi: 10.3389/fninf.2017.00076.

3. Johannes Köster and Sven Rahmann. Snakemake—a scalable bioinformatics workflow engine. Bioinformatics, 28(19):2520–2522, August 2012. ISSN 1367-4803. doi: 10.1093/bioinformatics/bts480.

4. Rebekah Loving, Delaney K. Sullivan, Fairlie Reese, Elisabeth Rebboah, Jasmine Sakr, Narges Rezaie, Heidi Y. Liang, Ghassan Filimban, Shimako Kawauchi, Conrad Oakes, Diane Trout, Brian A. Williams, Grant MacGregor, Barbara J. Wold, Ali Mortazavi, and Lior Pachter. Long-read sequencing transcriptome quantification with lr-kallisto. bioRxiv, July 2024. doi: 10.1101/2024.07.19.604364.

5. Ying Chen, Andre Sim, Yuk Kei Wan, Keith Yeo, Joseph Jing Xian Lee, Min Hao Ling, Michael I. Love, and Jonathan Göke. Context-aware transcript quantification from long-read RNA-seq data with Bambu. Nature Methods, 20(8):1187–1195, June 2023. ISSN 1548-7105. doi: 10.1038/s41592-023-01908-w.

6. Robert C Gentleman, Vincent J Carey, Douglas M Bates, Ben Bolstad, Marcel Dettling, Sandrine Dudoit, Byron Ellis, Laurent Gautier, Yongchao Ge, Jeff Gentry, Kurt Hornik, Torsten Hothorn, Wolfgang Huber, Stefano Iacus, Rafael Irizarry, Friedrich Leisch, Cheng Li, Martin Maechler, Anthony J Rossini, Gunther Sawitzki, Colin Smith, Gordon Smyth, Luke Tierney, Jean YH Yang, and Jianhua Zhang. Bioconductor: open software development for computational biology and bioinformatics. Genome Biology, 5(10):R80, 2004. ISSN 1465-6906. doi: 10.1186/gb-2004-5-10-r80.

7. Páll Melsted, Vasilis Ntranos, and Lior Pachter. The barcode, umi, set format and bustools. Bioinformatics, 35(21):4472–4473, May 2019. ISSN 1367-4811. doi: 10.1093/bioinformatics/btz279.

8. Josie Gleeson, Adrien Leger, Yair D J Prawer, Tracy A Lane, Paul J Harrison, Wilfried Haerty, and Michael B Clark. Accurate expression quantification from nanopore direct RNA sequencing with NanoCount. Nucleic Acids Research, 50(4):e19–e19, November 2021. ISSN 1362-4962. doi: 10.1093/nar/gkab1129.

9. Zahra Zare Jousheghani and Rob Patro. Oarfish: Enhanced probabilistic modeling leads to improved accuracy in long read transcriptome quantification. bioRxiv, March 2024. doi: 10.1101/2024.02.28.582591.

10. Hyun Joo Ji and Mihaela Pertea. Enhancing transcriptome expression quantification through accurate assignment of long RNA sequencing reads with TranSigner. bioRxiv, April 2024. doi: 10.1101/2024.04.13.589356.

11. Chen Yang, Justin Chu, René L Warren, and Inanç Birol. NanoSim: nanopore sequence read simulator based on statistical characterization. GigaScience, 6(4), February 2017. ISSN 2047-217X. doi: 10.1093/gigascience/gix010.

12. Rachael E. Workman, Alison D. Tang, Paul S. Tang, Miten Jain, John R. Tyson, Roham Razaghi, Philip C. Zuzarte, Timothy Gilpatrick, Alexander Payne, Joshua Quick, Norah Sadowski, Nadine Holmes, Jaqueline Goes de Jesus, Karen L. Jones, Cameron M. Soulette, Terrance P. Snutch, Nicholas Loman, Benedict Paten, Matthew Loose, Jared T. Simpson, Hugh E. Olsen, Angela N. Brooks, Mark Akeson, and Winston Timp. Nanopore native RNA sequencing of a human poly(a) transcriptome. Nature Methods, 16(12):1297–1305, November 2019. ISSN 1548-7105. doi: 10.1038/s41592-019-0617-2.

13. Nuala A. O’Leary, Mathew W. Wright, J. Rodney Brister, Stacy Ciufo, Diana Haddad, Rich McVeigh, Bhanu Rajput, Barbara Robbertse, Brian Smith-White, Danso Ako-Adjei, Alexander Astashyn, Azat Badretdin, Yiming Bao, Olga Blinkova, Vyacheslav Brover, Vyacheslav Chetvernin, Jinna Choi, Eric Cox, Olga Ermolaeva, Catherine M. Farrell, Tamara Goldfarb, Tripti Gupta, Daniel Haft, Eneida Hatcher, Wratko Hlavina, Vinita S. Joardar, Vamsi K. Kodali, Wenjun Li, Donna Maglott, Patrick Masterson, Kelly M. McGarvey, Michael R. Murphy, Kathleen O’Neill, Shashikant Pujar, Sanjida H. Rangwala, Daniel Rausch, Lillian D. Riddick, Conrad Schoch, Andrei Shkeda, Susan S. Storz, Hanzhen Sun, Francoise Thibaud-Nissen, Igor Tolstoy, Raymond E. Tully, Anjana R. Vatsan, Craig Wallin, David Webb, Wendy Wu, Melissa J. Landrum, Avi Kimchi, Tatiana Tatusova, Michael DiCuccio, Paul Kitts, Terence D. Murphy, and Kim D. Pruitt. Reference sequence (RefSeq) database at NCBI: current status, taxonomic expansion, and functional annotation. Nucleic Acids Research, 44(D1): D733–D745, November 2015. ISSN 1362-4962. doi: 10.1093/nar/gkv1189.

14. Andrey D. Prjibelski, Alla Mikheenko, Anoushka Joglekar, Alexander Smetanin, Julien Jarroux, Alla L. Lapidus, and Hagen U. Tilgner. Accurate isoform discovery with IsoQuant using long reads. Nature Biotechnology, 41(7):915–918, January 2023. ISSN 1546-1696. doi: 10.1038/s41587-022-01565-y.

15. Francisco J. Pardo-Palacios, Dingjie Wang, Fairlie Reese, Mark Diekhans, Sílvia Carbonell-Sala, Brian Williams, Jane E. Loveland, Maite D. María, Matthew S. Adams, Gabriela Balderrama-Gutierrez, Amit K. Behera, Jose M. Gonzalez Martinez, Toby Hunt, Julien Lagarde, Cindy E. Liang, Haoran Li, Marcus Jerryd Meade, David A. Moraga Amador, Andrey D. Prjibelski, Inanc Birol, Hamed Bostan, Ashley M. Brooks, Muhammed Hasan Çelik, Ying Chen, Mei R. M. Du, Colette Felton, Jonathan Göke, Saber Hafezqorani, Ralf Herwig, Hideya Kawaji, Joseph Lee, Jian-Liang Li, Matthias Lienhard, Alla Mikheenko, Dennis Mulligan, Ka Ming Nip, Mihaela Pertea, Matthew E. Ritchie, Andre D. Sim, Alison D. Tang, Yuk Kei Wan, Changqing Wang, Brandon Y. Wong, Chen Yang, If Barnes, Andrew E. Berry, Salvador Capella-Gutierrez, Alyssa Cousineau, Namrita Dhillon, Jose M. Fernandez-Gonzalez, Luis Ferrández-Peral, Natàlia Garcia-Reyero, Stefan Götz, Carles Hernández-Ferrer, Liudmyla Kondratova, Tianyuan Liu, Alessandra Martinez-Martin, Carlos Menor, Jorge Mestre-Tomás, Jonathan M. Mudge, Nedka G. Panayotova, Alejandro Paniagua, Dmitry Repchevsky, Xingjie Ren, Eric Rouchka, Brandon Saint-John, Enrique Sapena, Leon Sheynkman, Melissa Laird Smith, Marie-Marthe Suner, Hazuki Takahashi, Ingrid A. Youngworth, Piero Carninci, Nancy D. Denslow, Roderic Guigó, Margaret E. Hunter, Rene Maehr, Yin Shen, Hagen U. Tilgner, Barbara J. Wold, Christopher Vollmers, Adam Frankish, Kin Fai Au, Gloria M. Sheynkman, Ali Mortazavi, Ana Conesa, and Angela N. Brooks. Systematic assessment of long-read rna-seq methods for transcript identification and quantification. Nature Methods, 21(7):1349–1363, June 2024. ISSN 1548-7105. doi: 10.1038/s41592-024-02298-3.

16. Yukiteru Ono, Kiyoshi Asai, and Michiaki Hamada. Pbsim: PacBio reads simulator—toward accurate genome assembly. Bioinformatics, 29(1):119–121, November 2012. ISSN 1367-4803. doi: 10.1093/bioinformatics/bts649.

17. Dimitra Sarantopoulou, Thomas G. Brooks, Soumyashant Nayak, Antonijo Mrčela, Nicholas F. Lahens, and Gregory R. Grant. Comparative evaluation of full-length isoform quantification from RNA-Seq. BMC Bioinformatics, 22(1), May 2021. ISSN 1471-2105. doi: 10.1186/s12859-021-04198-1.

18. Heng Li. Minimap2: pairwise alignment for nucleotide sequences. Bioinformatics, 34(18): 3094–3100, May 2018. ISSN 1367-4811. doi: 10.1093/bioinformatics/bty191.

19. Simon A Hardwick, Wendy Y Chen, Ted Wong, Ira W Deveson, James Blackburn, Stacey B Andersen, Lars K Nielsen, John S Mattick, and Tim R Mercer. Spliced synthetic genes as internal controls in RNA sequencing experiments. Nature Methods, 13(9):792–798, August 2016. ISSN 1548-7105. doi: 10.1038/nmeth.3958.

